# Dentate gyrus mossy cells exhibit sparse coding via adaptive spike threshold dynamics

**DOI:** 10.1101/2022.03.07.483263

**Authors:** Anh-Tuan Trinh, Mauricio Girardi-Schappo, Jean-Claude Béïque, André Longtin, Leonard Maler

**Affiliations:** Kavli Institute for Systems Neuroscience, Norwegian University of Science and Technology, Trondheim, Norway, 7030; Department of Cellular and Molecular Medicine, University of Ottawa, Ottawa, Ontario, Canada, K1H 8M5; Department of Physics, University of Ottawa, Ottawa, Ontario, Canada, K1N 6N5; Brain and Mind Institute, Center for Neural Dynamics, University of Ottawa, Ottawa, Ontario, Canada, K1H 8M5; Center for Neural Dynamics, University of Ottawa, Ottawa, Ontario, Canada, K1H 8M5

## Abstract

Hilar mossy cells (hMCs) are glutamatergic neurons in the dentate gyrus (DG) that receive inputs primarily from DG granule cells (GCs), CA3 pyramidal cells and local inhibitory interneurons. The hMCs then provide direct excitatory and disynaptic inhibitory feedback input to GCs. Behavioral and *in vivo* single unit recording experiments have implicated hMCs in pattern separation as well as is in spatial navigation and learning. It has, however, been difficult to mechanistically link the *in vivo* physiological behavior of hMCs with their intrinsic excitability properties that convert their synaptic inputs into spiking output. Here, we carried out electrophysiological recordings from the main cell types in the DG and found that hMCs displayed a highly adaptive threshold acting over a remarkably protracted time-scale. The hMC spike threshold increased linearly with increasing current stimulation and saturated at high current intensities. This threshold also increased in response to spiking and this effect also decayed over a long timescale, allowing for activity-dependent summation that limited hMC firing rates. This mechanism operates in parallel with a prominent medium after-hyperpolarizing potential (AHP) generated by the small conductance K^+^ channel. Based on experimentally derived parameters, we developed a phenomenological exponential integrate-and-fire model that closely mimics the hMC adaptive threshold. This lightweight model is amenable to its incorporation into large network models of the DG that will be conducive to deepen our understanding of the neural bases of pattern separation, spatial learning and navigation in the hippocampus.

**Statement of significance:** Recent studies on hilar mossy cells have revealed that they are implicated in spatial navigation and mnemonic functions. Yet, the basic intrinsic characterization of these hMCs is still too superficial to explain their spiking behavior *in vivo*. Here, we describe novel biophysical properties of hMCs, including an independent relationship between spike latency and spike threshold as well as a slowly adapting spike threshold. These findings complement several other biophysical and connectivity similarities between hMCs and CA3 pyramidal cells, while emphasizing the contrast with hilar interneurons. Additionally, our results are well captured by a phenomenological model of the hMC which provides a useful framework to study the neural substrate of spatial navigation and learning in the dentate gyrus.

## Introduction

The dentate gyrus (DG) has long been discussed in relation to its putative roles in pattern separation and the generation of spatial maps (Marr, 1971; Leutgeb et al., 2007; Yassa and Stark, 2011; Neunuebel and Knierim, 2014). Given their large numbers, sparse activity and connectivity features (Diamantaki et al., 2016), DG granule cells (GCs) have been associated with pattern separation of their entorhinal cortex (EC) inputs (McNaughton and Morris, 1987; Rolls et al., 1998; Yassa et al., 2011). A subset of GCs are endowed with single place fields (GoodSmith et al., 2017), directly implying a role in spatial location encoding. These studies have rarely considered the excitatory hilar mossy cells (hMC) which receive GC input and then feedback to GCs (Scharfman, 1994a, 2016). Recent studies have however begun to directly implicate hMCs in both pattern separation (Jinde et al., 2012; Bui et al., 2018) and the encoding of spatial location (Danielson et al., 2017; GoodSmith et al., 2017; Senzai and Buzsaki, 2017). However, unlike their input GCs, single hMCs can discharge at multiple locations in space, thereby encoding multiple place fields (GoodSmith et al., 2017) and highlighting the singular properties of this back projecting pathway. Remarkably, neurons displaying connectivity (Elliott et al., 2017) and functional features (Fotowat et al., 2019) analogous to those of hMCs were recently described in a teleost analog of the hippocampal network, suggesting that hMC circuitry subserves a conserved role in the encoding of spatial maps.

The hMCs receive synaptic input from CA3 pyramidal cells as well as from GCs and the dynamic properties of these inputs onto hMCs have been studied (Scharfman et al., 1990; Scharfman, 1991, 1994a). The hMC feedback projections to GCs are both direct (excitatory) and via inhibitory interneurons (Scharfman, 1995) and the dynamic properties of these synapses have also been investigated (Hashimotodani et al., 2017; Hedrick et al., 2017; Lituma et al., 2021). These connectivity and physiology features suggest that hMCs act as a comparator of GC and CA3 activity (Elliott et al., 2017) that, via feedback, dynamically regulate the activity of the GC population. There is however limited information on the intrinsic biophysical properties of hMCs (Scharfman, 1992; Buckmaster et al., 1993; Scharfman, 1993; Buckmaster and Schwartzkroin, 1995), and this comprises obligatory knowledge to understand how this putative “comparator” computation naturally emerges from synaptic inputs. For example, *in vivo* recordings of hMCs have revealed that they often spike in a complex patterned manner (GoodSmith et al., 2017; Senzai and Buzsaki, 2017), but it unknown whether their spike output is simply a reflection of their dynamic synaptic inputs or whether intrinsic hMC dynamics also contribute to the patterning. Here, we present a combined electrophysiological and computational analysis that collectively identified salient features of intrinsic properties of hMCs, notably spike frequency adaptation mechanisms with a special emphasis on dynamic spike threshold. We compared the dynamic threshold across major cell types in the hippocampus: GCs, CA3 pyramidal cells, hilar interneurons and CA1 pyramidal cells to reveal the unique dynamics of hMCs, under the assumption that this will be most informative as to the functions of these “enigmatic” cells (Scharfman, 2016). The main features of this electrophysiological characterization were used to develop and validate a light-weight phenomenological exponential integrate-and-fire (EIF) model of hMC dynamics which can be implemented in a large-scale dentate gyrus network model to gain insights into of how they might integrate GC and CA3 pyramidal cell inputs to generate the spike patterns observed *in vivo*.

## Materials and Methods

For this study, we used 26-55 day old mice from the following transgenic mouse lines for cell type-specific identification; PV:TdTomato was obtained from crossing the PV-cre mouseline (B6.Cg-Pvalbtm4.1(flop)Hze/J; Jackson Labs; Stock No: 022730) with the Rosa:TdTomato (B6.Tg-Gt(ROSA)26Sortm14(CAG-tdTomato)Hze/J; Jackson Labs; Stock No: 007914) and the SOM:TdTomato was obtained from crossing the aforementioned RosaTdTomato mice with the SOM-cre (Sst-IRES-Cre; Jackson Labs; Stock No: 013044) mice. These mice were used to obtain recordings from hilar parvalbumin (PV) and hilar somatostatin (SOM) interneurons respectively. The hMC recordings were obtained from Drd2:TdTomato mice. The Drd2-cre (B6.FVB(Cg)Tg(Drd2-cre)ER44Gsat/Mmucd, MMRRC repository, Stock No: 032108-UCD) was crossed with the Rosa:TdTomato mouseline. For the other readily identifiable excitatory cell types (DG granule cells, CA3 pyramidal neurons and CA1 pyramidal neurons), recordings were obtained in wild type animals (C57BL/6 mice). Animals from both sexes were used and had access to food and water ad libitum. Rodents in this age range are capable of learning and our conclusions and model will therefore be applicable to studies of pattern separation and spatial learning in mice (Langston et al., 2010; Wills et al., 2010). All procedures were approved by the University of Ottawa Animal Care Committee and follow the guidelines from the Society for Neuroscience.

### *In vitro* slice procedure

Early experiments instructed us that we could reliably obtain quality recordings from hMCs neurons in slices prepared from young mice (<30 days old) using a slicing procedure routinely used for mice hippocampal recordings (Lee et al., 2016). However, hMCs in slices from older animals proved to be highly vulnerable and were not amenable to stable and long-lasting electrophysiological recordings. We therefore iteratively developed a slicing procedure to maximize hMC health for recordings in older animals. Prior to the dissection, an N-methyl-D-glucamine (NMDG) cutting solution and a modified recovery Ringer solution were placed in two distinct slice chambers. The NMDG solution was adapted from (Ting et al., 2014), and contained (in mM): 92 NMDG, 2.5 KCl, 1.25 NaH2PO4, 30 NaHCO3, 20 HEPES, 10 MgSO4, 25 Glucose, 0.5 CaCl2.2H2O, 5 Ascorbic Acid, 2 Thiourea, 10 N-acetyl-L-cysteine, 3 Sodium Pyruvate, and was adjusted to 295 mOsm while the modified recovery ACSF solution contained (in mM): 92 NaCl, 2.5 KCl, 1.25 NaH2PO4, 30 NaHCO3, 2 MgSO4, 25 Glucose, 2 CaCl2, 5 Ascorbic Acid, 2 Thiourea, 10 N-acetyl-L-cysteine, 3 Sodium Pyruvate, and was adjusted to 295 mOsm. Both recovery chambers were oxygenated (95% O_2_, 5% CO_2_) and heated to 37°C.

At the same time, another beaker containing the NMDG cutting solution (roughly 350 mL) was oxygenated and chilled to 4°C. Once the NMDG cutting solution was ready, the mouse was anesthetized by Isoflurane inhalation (Baxter Corporation, Canada). A transcardiac perfusion (10 mL) was then performed in order to exchange the animal’s blood with 3 mL of the chilled NMDG cutting solution after which the animal was sacrificed by decapitation. Once the brain was removed and placed in the chilled cutting solution, coronal sections (300 μm thick) were obtained using a vibratome (Leica) and the slices were transferred to the previously heated (to 37°C) custom-made incubation chamber containing the NMDG cutting solution. Before the slices were transferred to the heated NMDG cutting solution, both incubation chambers (one containing the NMDG cutting solution and the other, the recovery ACSF) were removed from the heating bath and were left to rest at room temperature. After 7-10 mins, the coronal slices were transferred to the other incubation chamber containing the recovery Ringer solution for at least 45 mins until the time of recording.

### *In vitro* recordings

Whole-cell recordings were carried out in recording ACSF (119 mM NaCl, 26 mM NaHCO3, 11 mM glucose, 2.5 mM KCl, 1 mM NaH_2_HPO_4_-H_2_O, 2.5 mM CaCl_2_, 1.35 mM MgSO_4_, and 295 mOsm, pH 7.4) in a perfused recording chamber at room temperature (23-25°C). Borosilicate glass micropipettes (Sutter Instruments) with resistances ranging between 5-12 MΩ, filled with a K-gluconate-based intracellular solution (135 mM K-gluconate, 7 mM KCl, 10 mM HEPES, 4 mM Mg-ATP, 10 mM phosphocreatine, and 0.4 mM Na-GTP, with an osmolality of 295 mOsm, pH 7.2), were used for recordings. Neurons were visualized with differential interference contrast (DIC) optics using a CMOS infrared camera (Scientifica). The recordings were first amplified using a Multiclamp 700B (Axon Instruments), filtered at 3 kHz and sampled at 10 kHz using a Digidata 1550 (Molecular devices). All recordings were carried out in current-clamp mode. At times, small DC current were applied to maintain the cell hyperpolarized at around −75mV. After attaining the whole-cell configuration, the resting membrane potential (RMP) was recorded for 30-60 s after which 500 ms square pulse currents of varying magnitude were injected into the cell. To characterize the slowly adapting spike threshold we encountered (see Results), we administered a ramp current injection protocol as previously described in (Trinh et al., 2019). Recorded cells were held for roughly 20-45 mins depending on the health of the cell. We generally limited current injection to a maximum of 200 pA to assure the stable long-term health of the recorded cells. In the case of hMCs we extended this current range to 300 pA in order to capture the full nonlinear response of these cells in our model (Fig. 3A, insert).

### Pharmacology

During this study, we used a glutamatergic receptor antagonist (10 μM cyanquixaline, or CNQX, Millipore-Sigma) and a GABAA chloride channel blocker (10 mM picrotoxin, or PTX, Abcam) to abolish synaptic activity during our recordings. Furthermore, we also used a Ca^2+^-activated K^+^ (SK) channel blocker, 30 μM UCL1684 (Tocris) to study the after-hyperpolarizing potential (AHP) in hMCs.

### Data analysis

Recordings were first visualized in Clampfit (Molecular devices) and then subsequently analyzed in Matlab (Mathworks) using custom scripts. To identify postsynaptic potential events, we developed a threshold-based method where the first derivative of the RMP was first smoothed with a moving average filter. Afterwards, an arbitrary threshold (0.05 mV*ms) was used to detect the peak of each event. The amplitude was calculated by subtracting the peak of each event with the baseline prior to each event. Given that this was a crude method for detecting events, we did not differentiate between individual and summed events which may have caused us to underestimate the true total of postsynaptic events. The spike threshold was defined as the value corresponding to an arbitrary chosen fraction (0.033) of the peak of the 1^st^ derivative of the membrane potential (Azouz and Gray, 2000; Trinh et al., 2019). The difference in spike threshold (or delta spike threshold) was defined as the difference between the first and nth spike threshold. Similarly, the delta spike height was calculated in a similar manner (difference between the heights of the 1st and nth evoked spikes). The latency to 1st spike was measured as the time from the onset of the intracellular current injection to the peak of the 1st evoked spike. The rheobase was defined as the minimal amount of current needed to produce an action potential. The AHP maximum amplitude was calculated as the difference between the spike threshold of the prior spike (spike “n”) and the minimum recorded value during the inter-spike interval (ISI) between spike “n” and spike “n+1”, *i.e.*, at the AHP trough. The hMC would sometimes emit spike doublets (or triplets) immediately upon strong current injections, therefore obscuring the full-time course of the AHP. In these cases, we therefore opted to study the AHP of the 2^nd^ spike (or 3^rd^ spike if there was a triplet burst). We estimated the total amplitude of the medium AHP (mAHP) area as the area of the AHP between its trough and extending to the membrane potential equal to the spike threshold of the previous spike (Fig. 2B, inset). All error bars were determined using the standard error of the mean. Wherever applicable, the statistical significance was determined using either the two-sample *t* test, the paired *t* test, or the Kruskal-Wallis test where *p* < 0.05 is considered significant (see statistical table). All final figures were assembled in Adobe Illustrator CS6 (Adobe Systems, RRID: SCR_010279).

### A model for the Hilar Mossy Cell dynamics

We were interested in finding a minimal model that is able to capture a number of features of the hMC data, foremost the threshold increase and decay following spikes. Our model is an essential component for modelling the *in vivo* responses of hMCs as animals learn spatial information and navigate. Given the lack of detailed knowledge of the kinetics of all channels expressed by hMCs (see below) we opted for an abstract model. We investigated the standard leaky (LIF) and the exponential integrate-and-fire (EIF) models and found that the latter could be expanded and subsequently fitted to reproduce the data. We did this process iteratively: we added a feature, then simulated the improved model and compared the following features with experiments: spike-dependent threshold increase, input-dependent threshold of the first spike, threshold decay, AHP effect (the dependence of the minimum of the AHP on the spike threshold), and the latency to the first spike. The simplest model that was able to capture these five features is given by

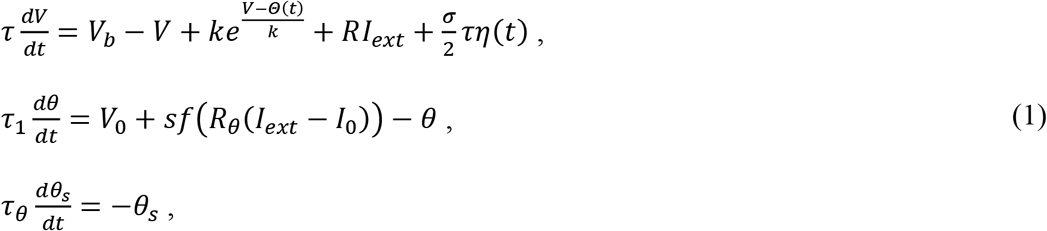

where *V* is the membrane potential, and the total threshold is dynamic and denoted by *Θ*(*t*) = *θ*(*t*) + *θ*_*s*_(*t*); *f*(*V*) = *V*/(1 + |*V*|) is a logistic function, and *η*(*t*) is a Gaussian white noise with zero mean and unit standard deviation that is included to generate variability in the model behavior; all the parameters are given in Table 1. In the standard EIF model, a spike occurs when the voltage runs off to infinity; a numerical criterion is then used to detect this upswing, followed by a manual resetting of the voltage. Accordingly, whenever *V*(*t*) > *V*_*peak*_, a >60 mV spike is said to have occurred. The voltage and the spike-dependent part of the threshold are then reset in the subsequent time step according to:

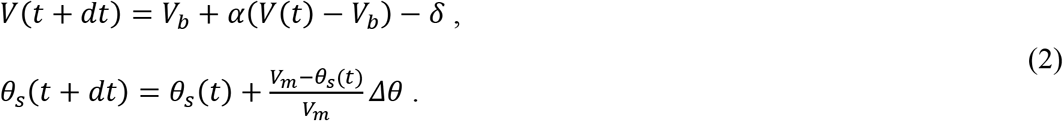

We did not need to include an absolute refractory period given the low firing rates and a mean action potential width of 5 ms; the voltage and threshold resets following a spike effectively mimic a refractory effect. We solve these equations using a second-order stochastic Runge-Kutta method (Honeycutt, 1992).

The exponential term in the voltage dynamics describes the rise of the spike more naturally as compared to a simple LIF (Fourcaud-Trocmé et al., 2003), *i.e.*, it mimics the positive feedback between voltage and sodium channel activation. Here, we replaced the standard EIF parameter for spike threshold *V*_*T*_ by a dynamic variable, *Θ*(*t*). This variable is composed of two terms: the input-dependent part, *θ*(*t*), is directly derived from experimental observations; it is counter-intuitive in that it increases as the external input increases, as observed in our data. It works similarly to an input-dependent activation gating variable, since the function *f*(*V*) is a sigmoid and plays the role of an activation function. We chose the logistic function instead of a Boltzmann function because of its numerical efficiency (Girardi-Schappo et al., 2017). The spike-dependent term *θ*_*s*_(*t*) in the total threshold saturates at *V*_*m*_ as described in Eq (2) (this term was introduced to enhance the fitting to the spike-dependent experimental data of hMCs), and it receives a discontinuous kick proportional to *Δθ* at every spike (Benda et al., 2010).

**Table 1.** Model parameters obtained by simulating the model and comparing the following features with experimental data: threshold increase versus spike number, threshold of the first spike versus input current, threshold decay, the covariance between AHP minimum and spike threshold, and the latency to the first spike. These values reflect an average hMC, generating curves that match the experimental observations within error bars. Finding a parameter set that matches a specific hMC would further require considering possible correlations between the parameters given here, and is beyond the scope of our work.

| Parameter | Description | Parameter | Description |
| --- | --- | --- | --- |
| $V_b = -67 \text{ mV}$ | Resting membrane potential | $k = 0.1 \text{ mV}$ | Spike slope |
| $R = 150 \text{ M}\Omega$ | Membrane resistance | $\tau = 38 \text{ ms}$ | Membrane time constant |
| $\sigma = 0.5 \text{ mV}$ | White noise standard deviation | $I_{ext}$ | External input |
| $\tau_1 = 20 \text{ ms}$ | Threshold activation time constant | $V_0 = -48.5 \text{ mV}$ | Threshold baseline |
| $s = 13.44 \text{ mV}$ | Threshold activation amplitude | $R_\theta = 20 \text{ M}\Omega$ | Threshold-input coupling resistance |
| $I_0 = 120 \text{ pA}$ | Threshold activation half current | $\Delta\theta = 2 \text{ mV}$ | Spike threshold increment |
| $V_m = 30 \text{ mV}$ | Spike-dependent threshold saturation | $\tau_\theta = 300 \text{ ms}$ | Threshold decay time |
| $\alpha = 0.8$ | Reset voltage slope | $\delta = 2 \text{ mV}$ | Reset voltage drop |
| $V_{peak} = -20 \text{ mV}$ | Threshold for resetting | | |

In the absence of noise, the model has a stable fixed point, corresponding to the resting membrane potential, as well as an unstable fixed point corresponding to a threshold. For a sufficiently large input current, the system goes into tonic spiking after an initial frequency adaptation due to the threshold and reset (AHP) dynamics. We measured the effective threshold of our EIF model by the same minimum derivative method described for the experiments. This is because the value of *Θ*(*t*) does not represent an actual hard threshold for spiking, even though it is connected to the unstable fixed point of the model. The nullcline dynamically shifts when the cell is stimulated due to the feedback between *Θ*(*t*) and the external input, such that the actual threshold becomes a function of time and has to be detected by the minimum derivative. In fact, the EIF model is known to present intrinsic threshold variability, and here we are introducing a new mechanism that causes the threshold to covary with both the injected current and the firing of the cell.

The reset rule of the membrane potential is also different from the standard EIF dynamics in Eq. (2). We assume that, after the spike, the membrane potential is directly proportional (by a factor less than one) to the potential right before the spike – a feature that is present in many neurons (Teeter et al., 2018). Here it is used specifically to implement a dependence of the AHP minimum on the spike threshold, compensating the post-spike threshold increase. Ultimately, this decreases the delay to the onset of the next spike, therefore increasing the evoked firing frequency when compared to a model with absolute reset and spike-dependent threshold. The membrane noise is added in order to mimic the statistical variability of averaging over multiple MCs for each of the considered features.

### Model simulation

We chose our model parameters so as to conform with the dependence of threshold and latency on input current. In addition, we attempted to simulate the dynamic threshold of hMCs since this is likely an important limiter of the stimulus-evoked spike rates *in vivo*. We simulated the step and ramp injection experiments (described in the *In vitro* recordings and Results sections) to probe for the threshold increase and decay. As in the experimental data, the threshold of the model was calculated by taking a fraction of the peak (0.033) of the first derivative of the membrane potential. The AHP minimum was estimated by taking the minimum value of the membrane voltage in between two consecutive spikes for a given injected current.

For the step simulations, we considered a wider range of inputs than in the experiments, since we wanted to probe for the general behavior of our modified EIF model. We considered currents from 70 pA to 400 pA. We ran 10 trials for each current and calculated the average and S.D. for each spike threshold and for the latency to the first spike. The threshold increase as a function of the spike number was calculated by further averaging the threshold values of each spike over all the injected currents in the range of 70 pA to 350 pA, yielding an average input 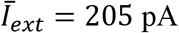. The threshold decay was calculated from the two-ramp injection simulation. We followed a protocol very similar to the one implemented experimentally (described in detail in the Results section). We first injected a stimulus ramp to excite the cell and cause the threshold to rise due to the spiking. Then, we inject a probe ramp after a certain delay to check the spike threshold at that later moment after the initial stimulation. We did 10 trials for each delay independently but kept the same maximum intensity and duration for both the stimulus (200 ms, 350 pA maximum) and the probe (100 ms, 300 pA maximum) ramps. The average threshold and S.D. were calculated for each delay.

### Code Accessibility

All codes related to the EIF model will be available upon request and they are also made available at the Github repository (https://github.com/mgirardis/mossy-cell-dg). A windows 10 computer was used to simulate the result from the EIF model.

## Results

Whole-cell electrophysiological recordings in current-clamp mode revealed that hMCs exhibit singularly large and frequent membrane potential fluctuations in the absence of any current injections (Fig. 1A). These fluctuations were often greater than 10-20 mV and at times generated spontaneous spiking: average rate of 0.48 ± 0.15 Hz (maximal observed rate of 1 Hz over 25 s) in a subset of our recordings where no holding current was used to artificially hyperpolarize the RM, inset Fig. 1A; N = 3). If we included cells that were hyperpolarized using a holding current, then 8/15 hMC cells displayed spontaneous spiking. These spontaneous spikes often occurred following the onset of large fluctuations, *i.e.*, >10 mV suggesting that multiple smaller coincident fluctuations are required to evoke spontaneous spiking (Fig. 1B). Previous studies have suggested that such fluctuations of the membrane potential may be caused by the stochastic opening and closing of various ion channels along the neuron’s membrane and producing an approximately Gaussian distribution of the mean-subtracted membrane potential (Faisal et al., 2008; Marcoux et al., 2016). The mean-subtracted membrane potential distribution observed in hMCs was asymmetrical (Fig. 1C). This suggests that the membrane noise may not be intrinsic in nature but instead driven by synaptic input.

**Figure 1.**
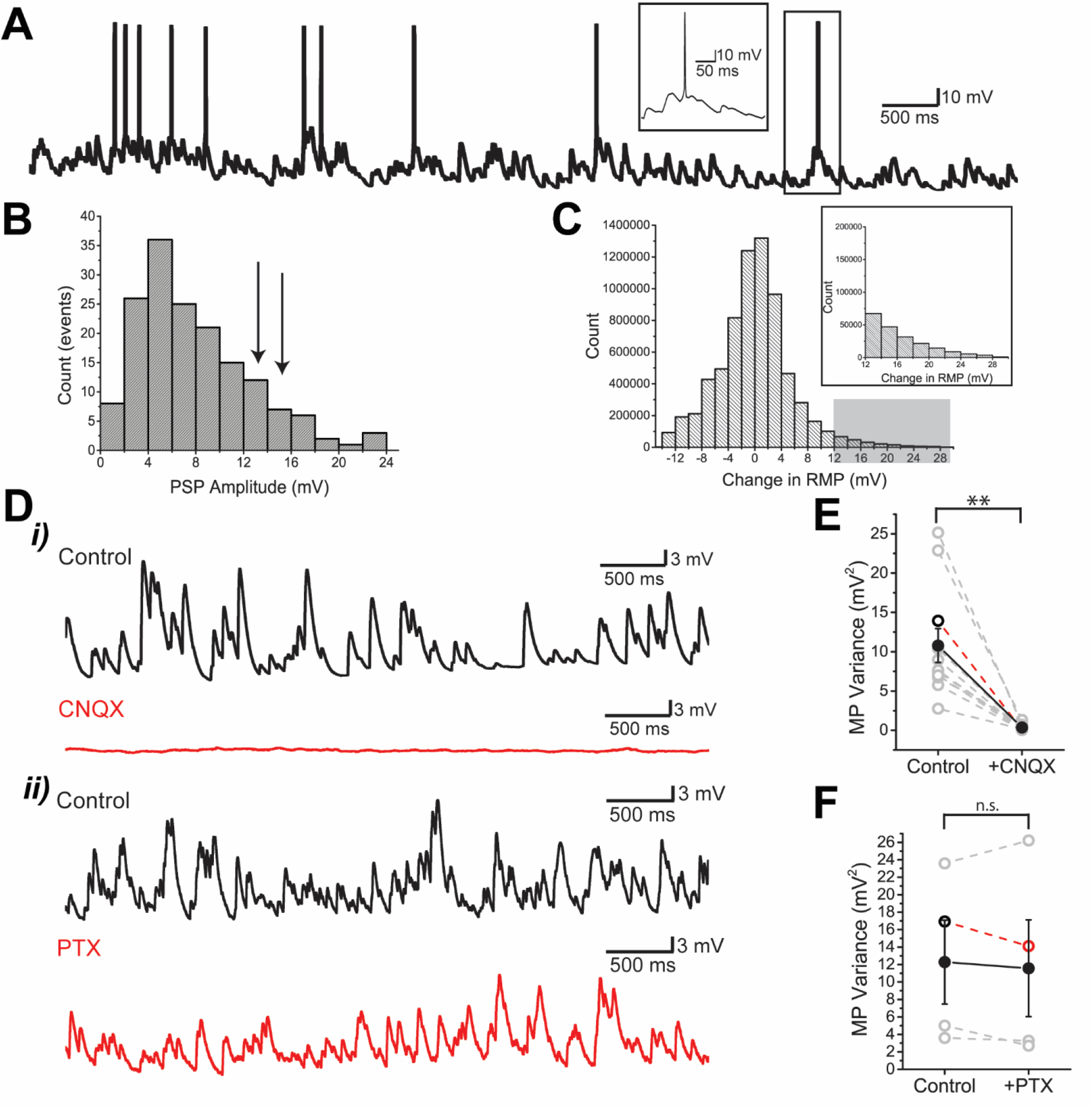
Strong membrane fluctuations are present in hMC neurons. **A.** Example of a cell displaying spontaneous spiking at a rate of 2.2 Hz. Multiple spontaneous events can be seen prior to spiking, which suggests that multiple coincident events are driving the cell to spike (inset). **B.** Postsynaptic potential amplitude histogram for the example trace shown in **A**. Analysis of the postsynaptic potentials reveals that most spikes occur during spontaneous events having greater than 12 mV in amplitude as denoted by the arrows. **C.** Average membrane fluctuations histogram. Each membrane potential value from all recording (N = 13 cells) were normalized by subtracting it with the mean of the membrane potential. The insert illustrates the long tail of the distribution which highlights the abundance of large events (>12 mV). **D.** Bath application of synaptic transmission blockers. **Di** The application of a glutamate AMPA receptor blocker, 10 μM CNQX, completely abolishes the generation of spontaneous events (bottom, red trace). **Dii** In contrast, the bath application of a GABAa antagonist, 0.1 mM PTX, did not appear to strongly affect the generation of spontaneous events. **E.** MP variance after the application of CNQX. The absence of synaptic events is apparent in the significant drop in MP variance. Each grey dashed line series represents an experiment, while each open circle represents the average for a given cell. The open circle black-red pair indicates the cell used in **1D**, while the filled black circle represents the average across all cells. **F.** MP variance after the application of PTX. The bath application of PTX did not significantly affect the MP variance. The same color scheme as in **1E** was used. n.s. = non-significant, ** p<0.01.

To test whether the observed noise is caused by spontaneous synaptic inputs, we bath applied the AMPA receptor antagonist CNQX (10 μM; N = 11 cells; Fig. 1Di). After this addition, the membrane potential variance dropped to negligible levels, from 10.8 ± 2.1 mV^2^ to 0.4 ± 0.1 mV^2^ (paired *t* test; p = 0.0005; row a, Table 2, Fig. 1E), demonstrating that these fluctuations were synaptically driven. Additionally, we quantified the contribution of GABA transmission membrane fluctuations by bath applying 0.1 mM picrotoxin (PTX), a GABAA receptor antagonist (Fig. 1Dii). The addition of PTX resulted in only a minor reduction of the membrane potential fluctuations (Control; 12.3 ± 4.8 mV^2^, PTX; 11.6 ± 5.5 mV^2^; paired *t* test; p = 0.61; row b, Table 2, Fig. 1F) in accordance with a previous study (Scharfman, 1993). This suggests that glutamatergic transmission was mostly responsible for the substantial fluctuations in hMC. Our results are consistent with earlier studies that have shown that the strong fluctuations of hMC membrane potential are due to spontaneously generated AMPA receptor mediated EPSPs; these studies have concluded that the likely source of these EPSPs is the DG granule cell mossy fiber projection to hMCs (Buckmaster et al., 1993; Scharfman, 1993). The presence of synaptic noise greatly hindered the estimation of hMC spike threshold and quantification of hMC AHPs. In the following experiments we therefore blocked both glutamatergic (CNQX) and GABAergic (picrotoxin) transmission in order to achieve clean recordings suitable for such measurements.

**Table 2.**
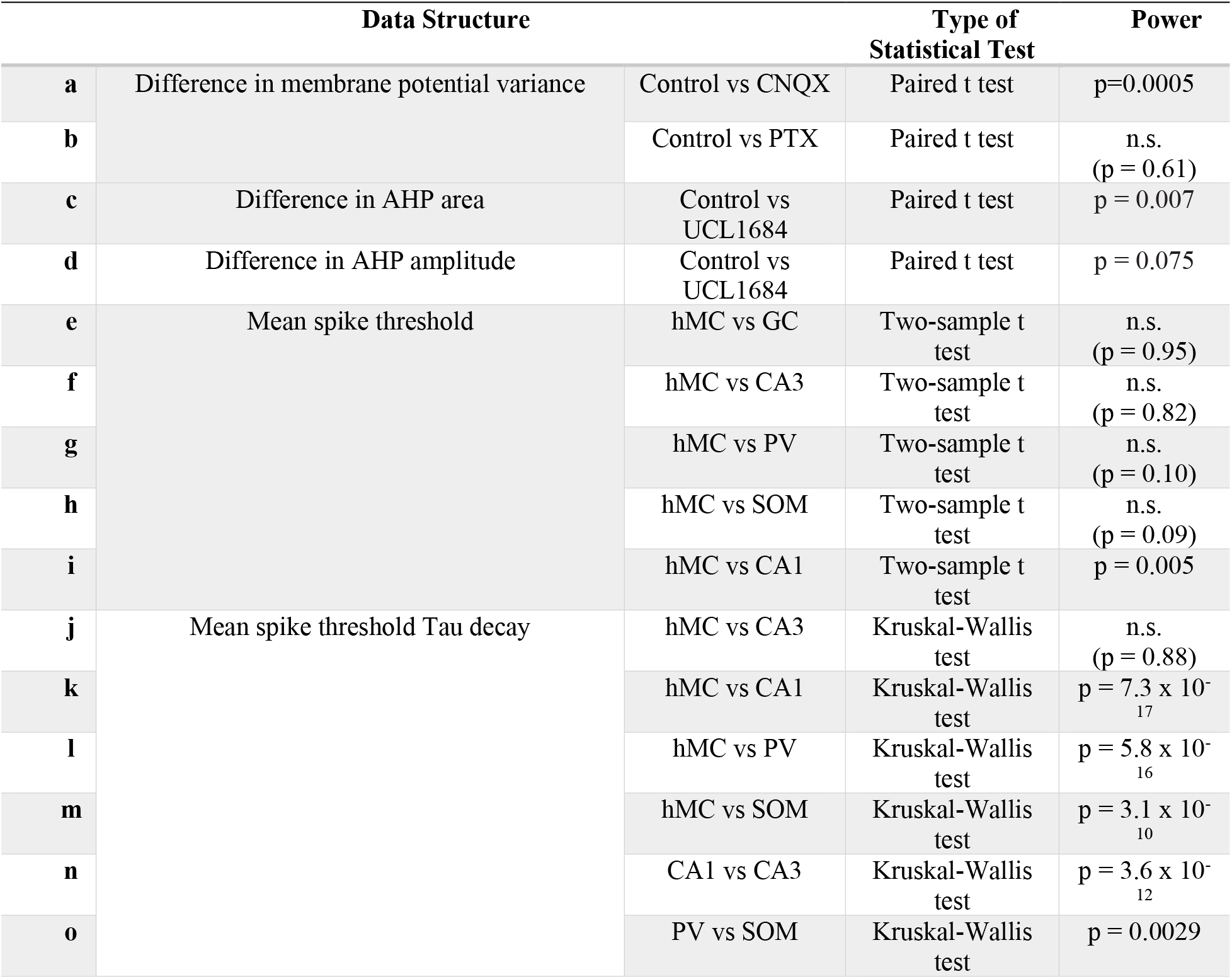
Statistical Table.

### Hilar Mossy Cell Intrinsic Properties

We next focused on the intrinsic biophysical properties of hMCs. To facilitate this analysis, these experiments were carried out in the presence of 0.1mM PTX and 10 μM CNQX. Using hyperpolarizing current injections, we measured a membrane time constant of 38.1 ±1.9 ms for the hMCs (used for the model below). Next, to examine the firing properties of these neurons, we injected square currents of various magnitudes. Remarkably, even strong current injection (+200 pA) hMCs did not induce high frequency spiking (average rate of 6.3 ± 1.1 Hz with +200 pA current injections, N =14 cells: Fig. 2Ai, Aii). We thus decided to examine the mechanisms that contribute to this low intrinsic excitability of hMCs.

**Figure 2.**
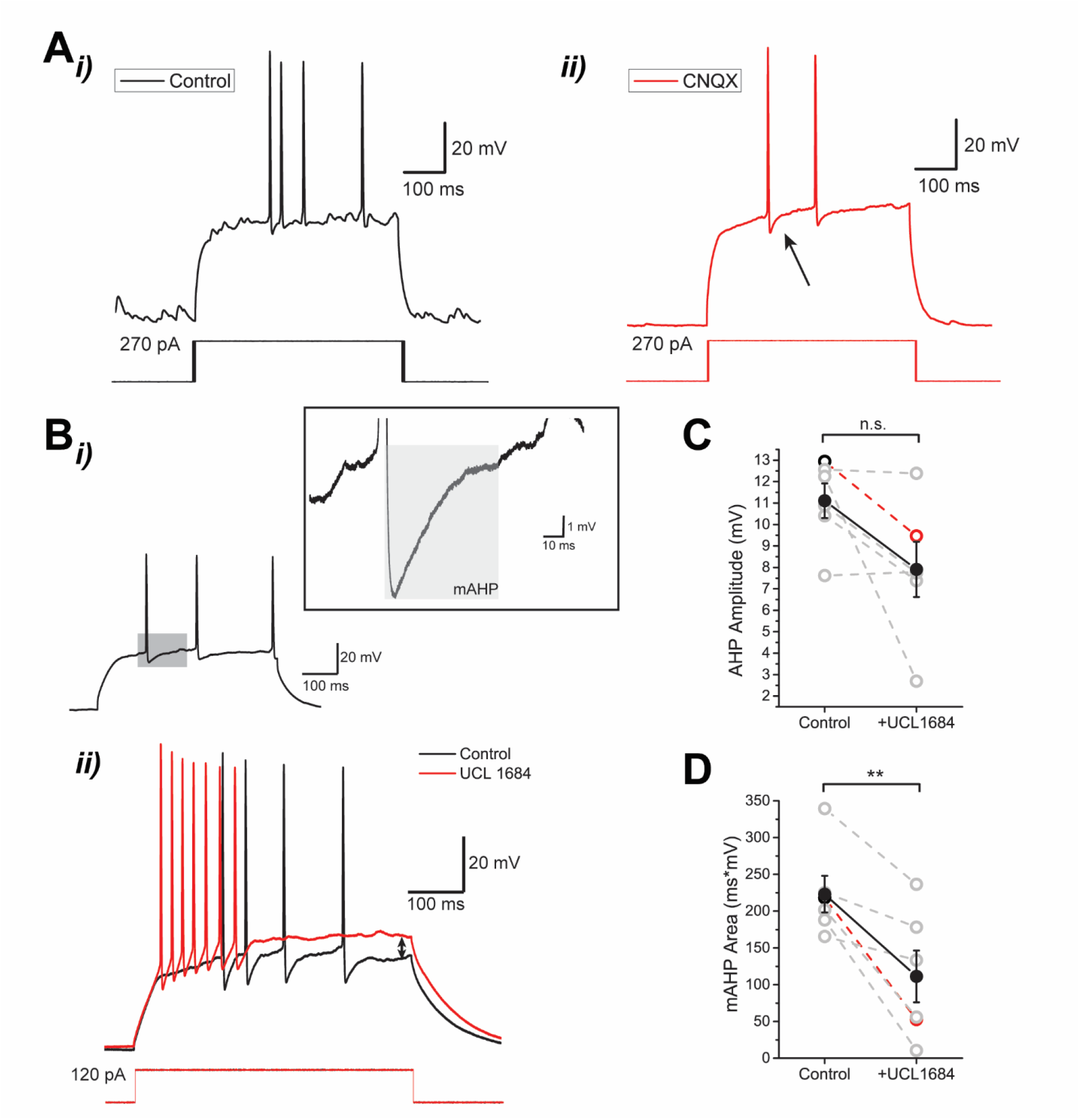
Characterizing the AHP in hMC neurons. **A.i)** Example neuronal response to +270 pA square pulse current injections. **ii)** Same as in **i)**, except that 10 μM CNQX has been applied prior to this recording. Notice that the AHP becomes more distinguishable in the absence of synaptic noise (black arrow). **Bi)** Left: Example response trace following a 500 ms square-pulse current injection in the presence of PTX and CNQX. A magnified view of the AHP delimited by the dark gray zone is shown in the inset on the right. Inset: The medium component of the AHP (mAHP) is defined by the light grey zone while the dark zone indicates the area under the curve of the illustrated AHP. The black arrow illustrates the trough of the AHP. **ii)** UCL1684 blocks SK channels and reduces the mAHP. In contrast, the recovery of the AHP was reduced as seen by aligning both traces. UCL also reduced the 1st spike threshold by ~3.0 mV and slightly depolarized the membrane potential as denoted by the black arrow at the end of the current response. **C.** mAHP area plot. In contrast to the amplitude, the area associated with the mAHP was significantly reduced by UCL1684 (N = 6 cells). **D.** AHP amplitude plot. The AHP amplitude was slightly reduced following the application of UCL1684, however, this was not significant (N = 6 cells). Each grey data point represents the average for a given cell. Each grey dashed line represents an experiment while each open circle represents the average for a given cell. The open circle black-red pair indicates the cell used in **Bii** while the filled black circle represents the average across all cells. n.s. = non-significant, ** p < 0.01.

Previous studies have shown that the medium AHP (mAHP) is driven by Ca^2+^-activated K^+^ channels, *i.e.*, the small conductance K^+^ (SK) channels (Kohler et al., 1996; Engel et al., 1999). Once the synaptic noise was removed by CNQX, we were able to visualize a prominent AHP which was previously masked by the barrage of PSPs (Fig. 2Aii, 2Bi). SK channels are expressed in neurons within the hilus of the rodent dentate gyrus (Cembrowski et al., 2016). Bath application of the SK channel blocker UCL1684 (30 μM) strongly affected the AHP (N = 6 cells; Fig. 2Bii) and induced a robust reduction in the area of the AHP (mean area for control: 223.1 ± 24.9; UCL1684: 111.3 ± 35.2 mV^2^, paired *t* test; p = 0.007; row c, Table 2, Fig. 2C). However, the peak magnitude of the AHP was only minimally affected by this antagonist (Fig. 2Bii; mean amplitude for control: 11.1 ± 0.8 mV; 7.9 ± 1.3 mV in UCL 1684; paired *t* test; p = 0.075; row d, Table 2, Fig. 2D). Thus, SK channels are a primary contributor of the mAHP in hMCs, although there are likely other channels that contribute to the fastest component of the AHP in these cells.

Bath administration of UCL1684 also caused hMCs to spike at membrane potentials that were previously subthreshold (Fig. 2Bii), suggesting that SK channels may be active at subthreshold potentials and slightly delay the firing of the 1^st^ spike. Previous studies have shown that SK channels are active at resting membrane potentials in mammalian neurons (Zhang and Huang, 2017). Furthermore, our previous study in the weakly electric fish (*Apteronotus leptorhynchus*) has also revealed this to be the case in pallial neurons (Trinh et al., 2019) but not in hindbrain electrosensory neurons (Ellis et al., 2007). Consistent with both these papers, we found that application of UCL1684 also caused the cell to fire more rapidly at first and then reach a more depolarized plateau potential (difference of ~9.2 mV between control and UCL1684 conditions; arrow in Fig. 2Bii). After the strong discharge, the hMCs cells stopped firing. We next examined the cellular basis of the strong spiking adaptation still present after the SK channel block.

Previous studies have shown that hilar mossy cells and hilar interneurons are highly interconnected, and both receive inputs from neighboring CA3 and DG granule cells (Amaral, 1978; Buckmaster and Schwartzkroin, 1994; Scharfman and Myers, 2012). We therefore compared intrinsic properties across these cell types because they could affect the reliability and temporal precision of the information transmitted by the hMCs within this network, notably spike threshold and latency to 1st spike. We have not injected currents larger than ~200 pA because most cells were not able to sustain spiking at higher current injections (>120 pA for GCs); further, we graph (Fig. 3) but do not report the analyses for GCs because of the limited current range we could use for these cells. We observed that the excitatory hippocampal neurons displayed linear increases in spike threshold in response to rising current injections (see Fig. 3A for linear fit values). The increase was shallow for CA1 pyramidal cells so that, over 70 to 180 pA current range, their threshold only increased from −54.0 ± 1.4 to −50.0 ± 0.7 mV. In contrast, CA3 pyramidal neurons and hMCs displayed steeper slopes (Fig. 3Ai) and, over the same current range, their threshold increased from −47.8 ± 3.2 to −38.2 ± 4.0 mV and from −50.4 ± 4.3 to −38.9 ± 2.7 mV respectively. Given our intention to model the hMC response to a wide range of synaptic currents, we stimulated them with currents up to 300 pA. As illustrated in Fig. 3Ai (inset), their spike threshold saturated at currents >200 pA. In sharp contrast, hilar interneurons did not show a stimulus intensity-dependent increase in spike threshold (Fig. 3Aii). A fixed, invariant threshold is what is typically expected and often modeled using the LIF formalism. Thus, these results outline a salient mechanism of hilar MCs and CA3 pyramidal wherein they adjust their spike threshold so that stronger excitatory synaptic inputs would presumably have to reach a higher threshold to initiate a spike (see Discussion).

**Figure 3.**
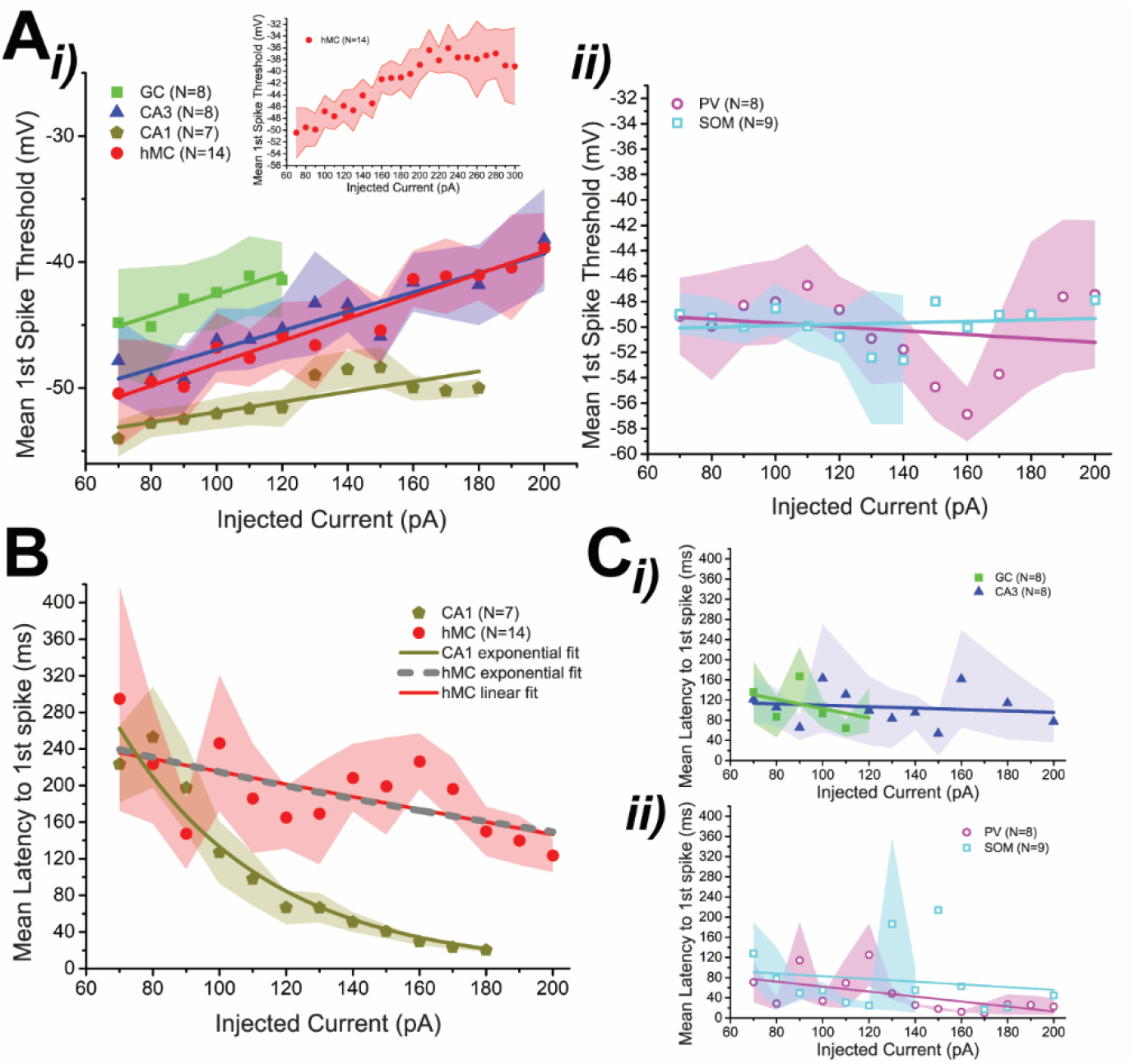
Stimulus dependent biophysical characteristics. **A.** Current-dependent increase in threshold for the first spike. **i)** Among the hippocampal excitatory cells, the CA1 pyramidal neurons showed the least amount of stimulus-dependent increase in spike threshold. In contrast, the excitatory cells within the hilar network all showed some degree of stimulus-dependent increase in spike threshold with the CA3 pyramidal neurons and the hMC showing similar dependencies (linear fits for CG: y = 0.084x - 50.98, CA3: y = 0.076x 54.59, hMC: y = 0.089x - 56.93, CA1: y = 0.04x - 55.93). Inset: At higher current injections, this increase in spike threshold saturates for the hMC. **ii)** In contrast, the hilar interneurons do not display this stimulus-dependent increase in threshold for the first spike (linear fits for PV: y = −0.015x - 48.17, SOM: y = 0.0057x 50.47). **B.** Current dependence of the latency to first spike. Stronger current injections cause the cell to spike earlier (hMC: N = 14 cells, CA1: N = 7 cells). An exponential fit was used for both hMC (dashed grey curve, y = 308.5\**e*^-x/276.1^, adjusted R^2^ = 0.33) and CA1 (dark yellow curve, y = 1279.3\**e*^-x/44.2^-25.6, adjusted R^2^ = 0.93). Given low correlation for the hMC, a linear fit was also used: red line, y = −0.69x + 284.16). **C.** Current dependency of the latency to first spike across other hippocampal neurons. In all cases, the stimulus intensity did not strongly affect the latency to first spike. **i)** Comparison across GC and CA3 pyramidal neurons. A linear fit was used to highlight the decrease in latency to first spike (GC: y = −0.92x + 195.36, CA3: y = −0.14x + 123.9). **ii)** Comparison of the hilar interneurons. A linear fit was used to highlight the decrease in latency (PV: y = −0.49x + 111.76, SOM: y = −0.27x + 109.94). Color scheme: GC: green, full squares, N = 8 cells; CA3: blue triangles, N = 8 cells; CA1: dark yellow pentagons, N = 7 cells; hMC: hilar MC: red circles, N = 14 cells; hilar PV: magenta, open circles, N = 8 cells; hilar SOM: open cyan squares, N = 9 cells. In all figures, the shaded area corresponds to the standard error of the mean.

For a neuron with a passive (RC) subthreshold membrane, a stronger stimulus current is expected to more rapidly charge the membrane capacitance resulting in a shorter first spike latency. In contrast, the increased spike threshold outlined above is expected to counteract this effect and increase latency. We therefore examined the latency to 1^st^ spike as a function of current intensity to clarify the effect of a dynamic threshold on spike latency. In CA1 pyramidal neurons, we found that increasing the stimulus current intensity caused a simple exponential decrease in spike latency (Fig. 3B; dark yellow curve): from 223.3 ± 41.9 to ~20.2 ± 1.9 ms (~200 ms; Fig. 3B), consistent with the relatively small increase in threshold observed over this current range. Hilar MCs spike latency in latency was best captured by a linear fit and latency decreased from 295.1 ± 122.7 to 123.9 ± 18.4 ms (~170 ms; Fig.3B). The greater increase in threshold for hMCs vs CA1 pyramidal cells may account for the smaller decrease in latency, but the very noisy hMC latency vs current plots suggests that other factors are operative. Surprisingly, CA3 pyramidal cell and interneuron spike latencies were nearly independent of current intensity (Fig. 3Ci, 3Cii).

Altogether, we conclude thus far that the dynamic spike threshold and latency can be at least partly independently controlled by unknown subthreshold mechanisms. We further note that the latency of hilar interneurons was far lower than that of hMCs, *i.e.*, a weak input to an interneuron will initiate spiking at a shorter latency than a strong input to a hMC (compare Fig. 3B vs 3Cii).

### Dynamic spike threshold in the hippocampal formation

Based on the analysis outlined above, it seems that the spike threshold is strongly regulated in hMCs and thus warrants further investigation. When examining the first evoked spike from square-pulse current injections, we observed that the distribution of 1^st^ spike thresholds in hMCs was quite similar whether CNQX was applied or not (N = 14 cells: Fig. 4Ai). Additionally, the spike threshold did not seem to correlate with the variability of the resting membrane potential (N = 14 cells: Fig. 4Aii). When we compared the average spike threshold across the major hippocampal cell types, we found that the average spike threshold of the hMC (mean 1^st^ spike threshold: 42.6 ± 2.0 mV) was comparable to the other major hippocampal cell types tested (GC, N= 8 cells: 42.3 ± 2.7 mV; CA3, N = 8 cells: −41.7 ± 2.9 mV; hilar PV, N = 8 cells: −47.7 ± 2.4 mV; hilar SOM, N = 9 cells: −48.0 ± 2.5 mV; two-sample *t* test: hMC vs GC; p = 0.95, row e, Table 2; hMC vs CA3; p = 0.82, row f, Table 2, hMC vs PV; p = 0.10, row g, Table 2; hMC vs SOM; p = 0.09, row h, Table 2; Fig. 4Aiii). The one exception was CA1 pyramidal neurons which had a significantly lower threshold (CA1, N = 7 cells, mean 1st spike threshold: −51.4 ± 1.4 mV; two sample *t* test; hMC vs CA1; p = 0.005, row i, Table 2; Fig. 4Aiii). This comparison reveals that that the excitatory cells within the hilar network have similar spike thresholds that are slightly lower than those of the inhibitory interneurons (not significant) and significantly lower compared to the CA1 pyramidal neurons.

**Figure 4.**
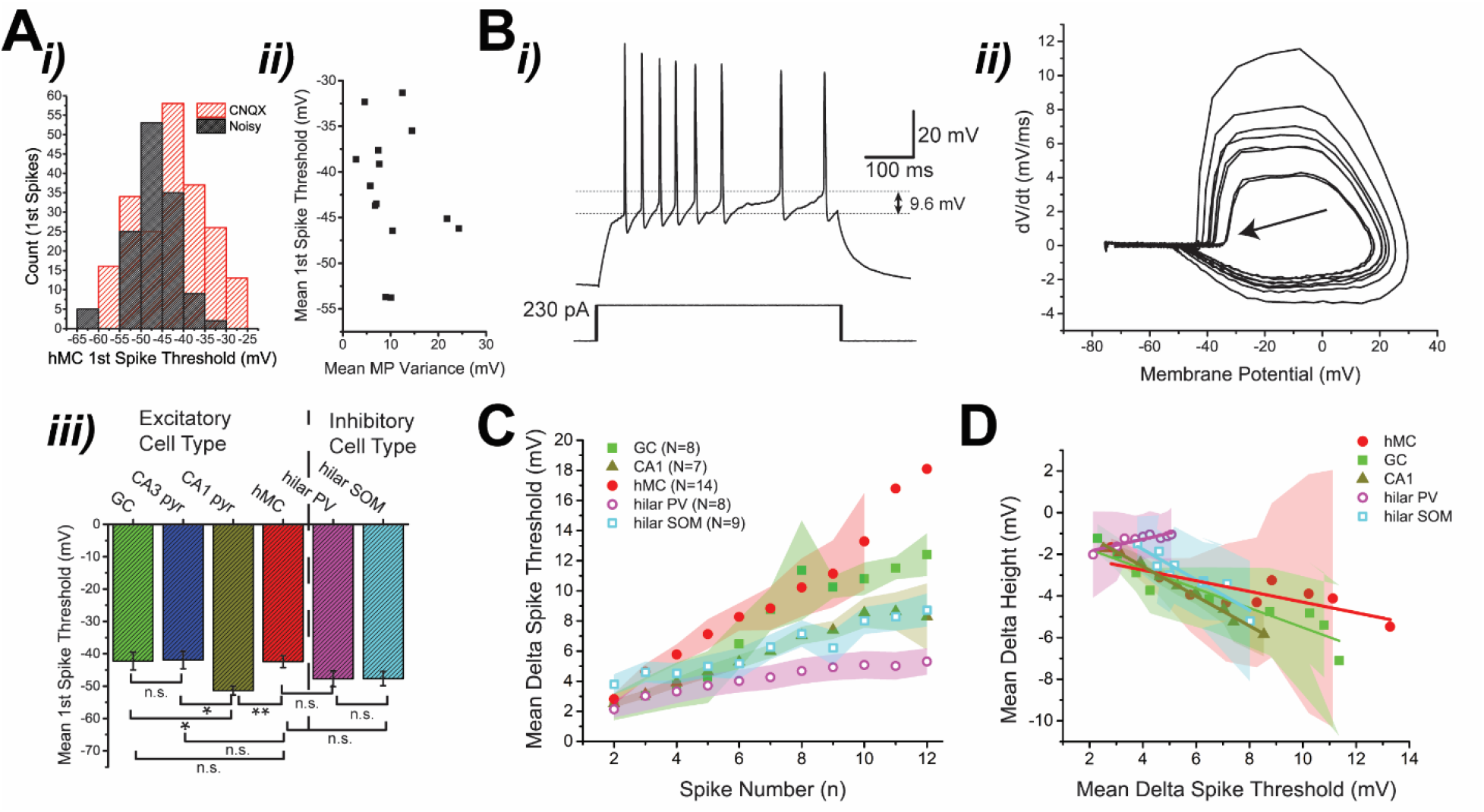
Comparison of the spike threshold dynamics in various hippocampal neurons. **A.i)** Histogram of the first spike threshold in the presence of synaptic noise (black) and in the absence of synaptic noise (red) in hMCs. **ii)** The effects of MP fluctuations to the spike threshold in hMCs. By comparing the mean first spike threshold with the mean MP variance, we observed that most cells with high spike threshold had low MP variance prior to the bath application of CNQX (N = 14 cells). Here each square represents the mean value for a given cell. **iii)** Mean first spike threshold comparison across different hippocampal cell types (GC: N = 8 cells, CA3: N = 8 cells, CA1: N = 7 cells, hMC: N = 14 cells, hilar PV: N = 8 cells, hilar SOM: N = 9 cells). **B.** The recorded spiking response displayed a dynamic spike threshold. **i)** This example response trace to a +230 pA current injection displayed an increasing spike threshold after successive spiking. When comparing the first spike to the last spike, there is a difference of 9.6 mV. **ii)** This effect can be further illustrated by the phase plot of the membrane potential during the current injection. The black arrow illustrates the last spike of the neuron’s response. **C.** Comparison of the dynamic spike threshold across hippocampal cell types. The delta spike threshold (see Methods) was plotted as a function of the spike number for all major hippocampal subtypes (red circle: hMC, green square: GC, dark yellow triangle: CA1; magenta open circle: hilar PV, cyan open square: hilar SOM). CA3 pyramidal neurons were not shown here due to the low numbers of recorded spikes. The shaded area corresponds to the standard error of the mean. **D.** Spike threshold adaptation and spike height adaptation. The increase in spike threshold (delta spike threshold) was plotted as a function of the delta height (difference in spike height; see Methods). A linear fit was used to highlight the decrease in spike height (GC: y = −0.47x - 0.84, CA1: y = −0.71x - 0.26, hMC: y = −0.26x - 1.74, PV: y = 0.29x - 2.43, SOM: y = −0.73x - 1.14). Same color scheme as in **4C**. n.s. = non-significant, * p<0.05, ** p<0.01.

Past *in vivo* studies have demonstrated that an hMCs often displayed complex spiking patterns when the experimental animal (rat, mouse) traversed its place field (Goodsmith et al., 2017; Senzai et al., 2017). This prompted to investigate whether there are other intrinsic mechanisms that could control the spiking behavior of these cells. In addition to the AHP, a second intrinsic mechanism which could regulate spiking is a dynamic spike threshold (Trinh et al., 2019). When examining the evoked spiking response, we noticed that the spike threshold increased after successive spiking (Fig. 4B). A slowly recovering spike threshold increase was found not only in hMCs but also in all the other studied hippocampal cell types (Fig. 4C). CA3 pyramidal neurons were omitted in this analysis, given their inability to produce sufficient spiking for a valid comparison across cells (data not shown). Despite being present in multiple cell types, this spike frequency adaptation mechanism was strongest in the hMCs when compared to the other hippocampal cell types, with the hilar interneurons showing the least amount of adaptation (Fig. 4C).

It has previously been shown that a dynamic spike threshold adaptation may be caused by the slow inactivation of Na^+^ channels which accumulates after successive spiking (Platkiewicz and Brette, 2011; Trinh et al., 2019). Such inactivation of Na^+^ channels can be measured indirectly as a reduction of spike height, since it is presumed that the inactivation will cause fewer Na^+^ channels to be opened by a depolarizing input (Henze and Buzsaki, 2001). Effectively, this increased in spike threshold (delta threshold) was shown to be negatively correlated with a reduction in spike height (delta height) which occurred after successive spiking in all measured cell types, except in PV interneurons (Fig. 4D). These results suggest that an accumulation of Na^+^ channel inactivation may also be responsible for the increased in spike threshold observed in these cells.

To further characterize the dynamic spike threshold recovery, we used a ramp current injection protocol which had been previously used to measure the decay of this adaptation process in hippocampal-like cells of a teleost fish (Trinh et al., 2019). By injecting a long ramp current to evoke spiking and threshold fatigue (stimulus ramp), followed by a shorter ramp current injection (probe ramp) at various time intervals, we measured the change in spike threshold as a function of inter-stimulus interval duration (Fig. 5Ai, 5Aii). In the case of the hMCs, the increase in spike threshold accumulated at the end of the protocol and relaxed back to its resting state by the end of the protocol (Fig. 5Aiii). Surprisingly, the recovery from this spike threshold fatigue in hMCs followed a similar timescale to the fish hippocampal-like neurons (mean decay time constant = 570 ± 70 ms; Fig. 5B).

**Figure 5.**
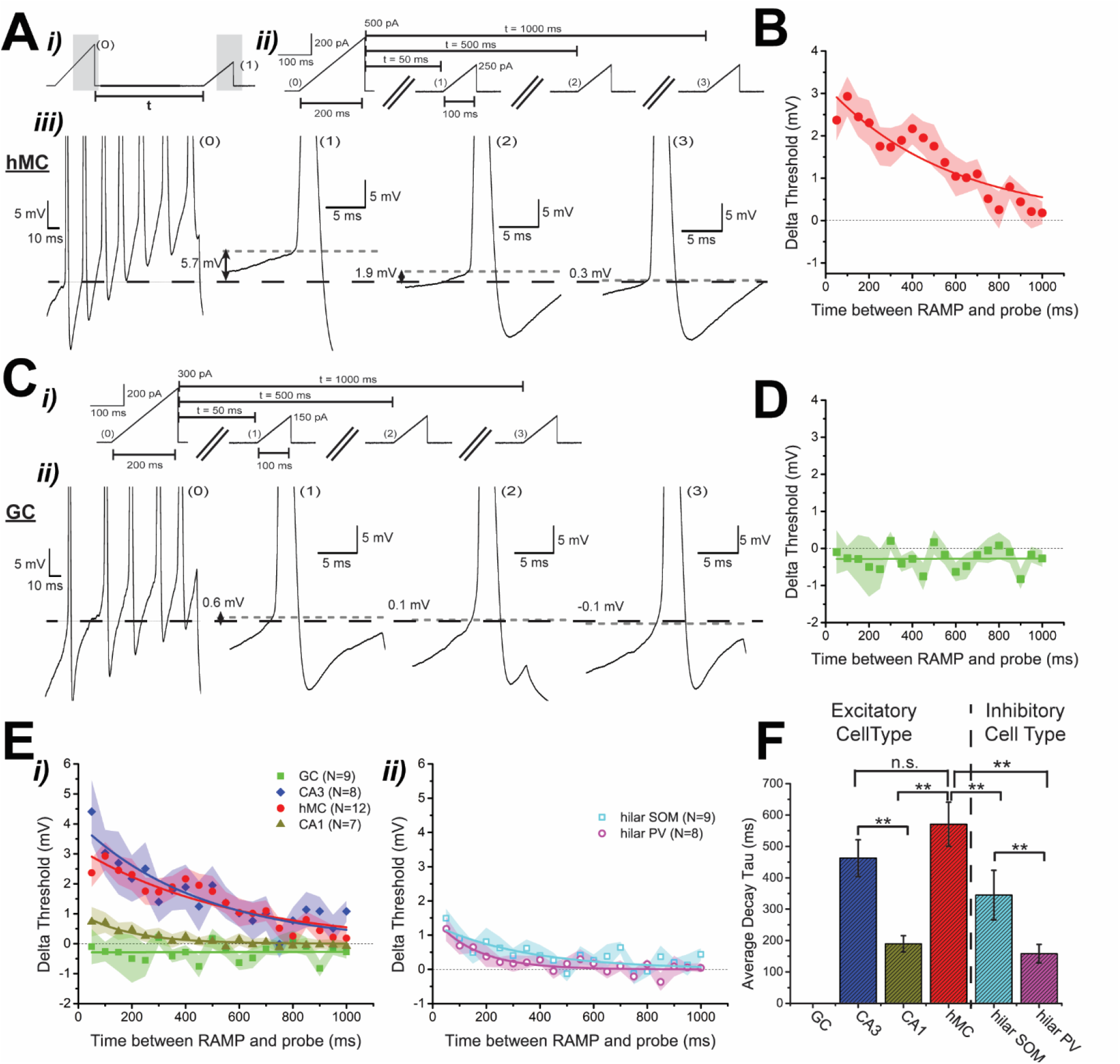
Spike threshold adaptation timescale. **A.** Example hMC response to the ramp current injection protocol. **i)** Two ramp currents (0) and (1) separated by an adjustable time t were injected **ii)** The first stimulus ramp current (0) was always twice as strong and twice as long in duration as the subsequent probe ramp (1). In this example the stimulus ramp current was +500 pA and lasted 200 ms while the probe ramp current was +250 pA and lasted 100ms. By injecting the probe ramp current at different pre-selected times *t* (in this example, 50, 500, 1000ms), we can measure the recovery from the spike threshold adaptation. **iii)** Example hMC response to the stimulus ramp current injection (0), and various probe ramp current injections after *t* = 50 ms (1), *t* = 500 ms (2), *t* = 1000 ms (3). When the time *t* between the two ramp current injections was short, a strong increase in spike threshold was observed (1), while when the time *t* was long (3), the increase in spike threshold was negligible. **B.** Decay of the spike threshold adaptation in hMC. The difference in spike threshold between the first spike resulting from the stimulus ramp current injection and the probe ramp current injection was plotted as a function of time separation *t*. The decay curve was fitted with an exponential (y = 3.175*e*^-x/570.42^). The shaded area corresponds to the standard error of the mean. **C.** Example DG granule cell’s response to the ramp current injection protocol. **i)** The same protocol described in **5i)** was used, except the scale of the current injection was different (+300 pA instead of +500 pA). **ii)** Example GC response to the stimulus ramp current injection (0), and various probe ramp current injections after *t* = 50ms (1), *t* = 500ms (2), *t* = 1000ms (3). Unlike the hMC, the GC did not show any persistent changes in spike threshold that adapts over time. **D.** Same as in **5B**, except for DG granule cells. Since theses neurons do not show any prolonged spike threshold increase, a linear fit was used instead of an exponential fit (y = 0.0000139x - 0.287) **E.** Comparison of the spike threshold adaptation in hippocampal neurons. **i)** Same as in **5B** and **5D**, except that all the excitatory neuron subtypes are presented (red circles: hMC, green squares: GC). An exponential fit was also used for the CA3 (blue diamonds: y = 4.031*e*^-x/462.58^) and for the CA1 neurons (dark yellow triangles: y = 1.0432*e*^-x/189.4^) **ii)** Same as in **5B** and **5D**, except for all hilar interneurons. An exponential fit was also used for the hilar PV (magenta open circles: y = 1.521*e*^-x/157.76^) and for the hilar SOM (cyan open squares: y = 1.326*e*^-x/344.76^) neuron. **F.** Comparison of the average decay tau of the spike threshold adaptation in hippocampal neurons. n.s. = non-significant, ** p<0.01.

We next examined spike threshold adaption in the other cell types of the hippocampus. DG granule cells showed no significant spike threshold adaptation over long periods of time (Fig. 5C, 5D). CA3 pyramidal neurons and hMCs displayed the strongest increase in spike threshold (CA3 mean: 4.41 ± 1.05 mV, hMC mean: 2.38 ± 0.45 mV) 50 ms following the first ramp current injection. Both CA1 pyramidal neurons and the inhibitory cell types showed little increase in spike threshold (Fig. 5Ei, 5Eii). All hippocampal cell types, except for the DG granule cells, were fitted with an exponential curve which allowed us to estimate the average decay time constant of the spike threshold adaptation. Based on our fitting, the recovery from the spike threshold fatigue was slowest in the CA3 (mean tau: 463 ± 59 ms) and hMC (mean tau: 570 ± 70 ms; Fig. 5F) neurons when compared to the other cell types (mean tau: CA1: 189 ± 26 ms; hilar SOM: 345 ± 79 ms; hilar PV: 158 ± 30 ms; Kruskal-Wallis test: hMC vs CA3; p = 0.88, row j, Table 2; hMC vs CA1; p = 7.3 x 10^−17^, row k, Table 2, hMC vs PV; p = 5.8 x 10^−16^, row l, Table 2; hMC vs SOM; p = 3.1 x 10^−10^, row m, Table 2; CA1 vs CA3; p = 3.6 x 10^−12^, row n, Table 2; PV vs SOM; p = 0.0029, row o, Table 2). Considering the slow timescale of this dynamic spike threshold adaptation, we can therefore hypothesize that this process was caused by the slow recovery of Na^+^ channels inactivation as previously shown in the hippocampal-like neurons of the weakly electric fish (Trinh et al., 2019).

### An effective model for the threshold dynamics

Aiming at the future development of a hilar network model, we initially set out to develop a single cell model which captures salient excitability features of hMCs, notably their spike threshold dynamics (see Methods for model choice justification, and simulation details). During the fit process, we considered several variations of the IF model, and a few of them were good in fitting most of the experimental features in the hMCs. We also considered other quantities, like AHP amplitude and ISI while adjusting the model. We started from a leaky IF with a simple spike-dependent dynamic threshold (Benda et al., 2010). This base model is given in Eqs. (1)–(2) with the following simplifications: remove the exponential term in the membrane potential, replace the membrane reset expression by an absolute value *V*_*R*_, and replace the input-dependent dynamic *θ*(*t*) by a constant *θ*_0_ that represents the average initial threshold of hMCs. Then, the total threshold is *Θ*(*t*) = *θ*_0_ + *θ*_*s*_(*t*), and the cell emits a spike whenever *V*(*t*) = *Θ*(*t*). This model fitted the threshold increase versus spike number, but performed poorly for all other features, since there is no dependency on the intensity of the injected current, nor covariance of the AHP with the threshold.

The first modification that we tried on the leaky IF was to replace the average initial threshold by a variable *θ*(*t*) whose rate of change varied linearly with the membrane potential, *dθ*/*dt*~*V*. This model has an analytical solution, and it can can be shown that the first spike threshold decreases as the injected current increases (not illustrated), contrary to what is observed in hMCs. The observed dependence of the first spike threshold in Fig. 3A (inset) motivated the inclusion of a sigmoidal shape to 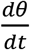 that depended on the external current, rather than on *V*, as described in Eq. (1). An improved leaky IF with dynamic threshold then included this feature together with the linear resetting rule for *V*, resulting in a model almost equal to the one given by Eqs. (1) and (2), except for the exponential term in the membrane equation. This improved linear IF also had an extra simplified K^+^-current for the AHP, *I*_*K*_ = −*g*_*K*_(*V* − *E*_*K*_), with *g*_*k*_ obeying an equation similar to *θ*_*s*_ (a positive kick on the reset, followed by an exponential decay). This version had a very nice overall fit to most of the parameters and performed similarly to our modified EIF that we showed in this section (see Fig. 6). However, the fit to both the threshold decay and to the first spike threshold could not be done simultaneously.

**Figure 6.**
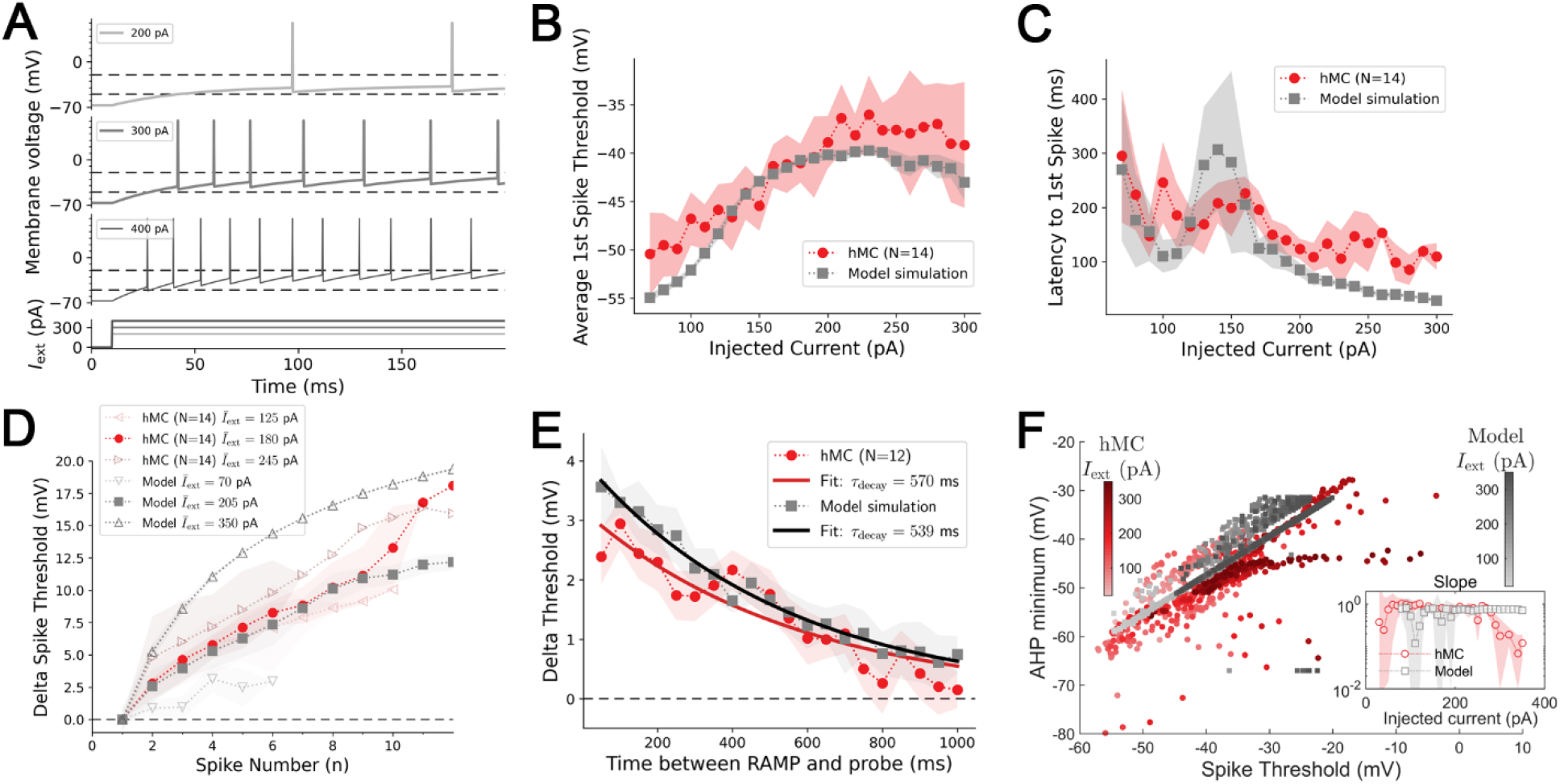
Model simulation. **A.** Representative spikes for three different step current intensities. This protocol is used to probe for the model properties in all the other panels (except for the threshold decay in panel **E**, where the ramp input protocol is used as explained in Fig. **5**. Dashed lines are guides to help identify the increase of the threshold and AHP depth. **B.** The input-dependent threshold follows closely the sigmoidal trend observed experimentally. **C.** The latency to first spike tends to diverge for small inputs similarly in model and experiments. **D.** Spike-dependent threshold increase; the experimental data (circles) are the same as presented in Fig. **3C** for hMCs. The side-pointing empty triangles are the experimental data averaged over high (190 to 300 pA) and low (70 to 180 pA) input currents. The model threshold increase was averaged over input current intensity (squares) ranging from 70 to 350 pA. Up- (down-)pointing triangles show the behavior of the model for high (low) injected current. **E.** The decay has a similar time constant and follows closely the one from hMCs. **F.** Covariance of the AHP minimum potential with the spike threshold, supporting the linear reset rule implemented in the model. The slope of the covariance remains constant at 0.8 for a wide range of inputs (inset) in both experiments and model (hence the chosen *α* in Table 1). All shaded areas are standard errors except for panel **F** inset, where it represents the 95% confidence interval from the best fitting slope of the covariance.

We then decided to explore the EIF, since it has a more natural spike initiation, without the need of an artificial threshold (Fourcaud-Trocmé et al., 2003). By introducing a novel dynamic into its threshold-related parameter, which controls the unstable fixed point of the underlying dynamics, we were able to reach a compromise between a good fit to both the threshold at spike initiation and the threshold decay. We were also able to drop the K^+^-current, which had been introduced for the leaky IF to enhance the AHP amplitude fitting. This is because the exponential dynamics adds more flexibility to the membrane potential equation and is capable of generating a good enough AHP without that explicit current.

The EIF fits to all the considered hMC features are detailed in Fig. 6. For example, the input-dependent part of the modified EIF threshold follows closely the hMC data, as expected: it presents a sigmoidal dependency on the injected current, saturating at around 200 pA (Fig. 6B). The latency, however, is underestimated for large currents, but matches the experimental data in the weakly excitable regime, when the injected current is very small (Fig. 6C). This is because hMCs tend to have a consistent delayed first spike, even though the delay slowly decreases with input intensity. The change of slope before the first spike in hMCs (as in the control condition of Fig. 2Bii), as well as the underestimation of the first spike latency, suggest that there is an underlying mechanism delaying the first spike. To account for this fact, we tried implementing a simplified fast inactivating K^+^-current, also known as an IA current, as proposed for serotonergic neurons in the raphe nucleus (Harkin et al., 2021) since these currents are also known to be present in mossy fiber boutons (Geiger and Jonas, 2000). However, the IA currents caused the rheobase currents in our model to rise significantly, such that we were able to initiate spiking only for currents greater than 150 pA, contrary to what we observed in hMCs (some cells could spiked with current injections as low as 20 pA, data not shown). This suggests that a deeper scrutiny of the hMC peri-threshold membrane currents is necessary to be able to fully capture the delay to first spike.

In hMCs, the spike-dependent threshold increase is more consistent than in the modified EIF model (Fig. 6D): for small inputs (mean 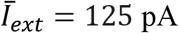), the spike-dependent threshold of hMCs grow almost as much as for large inputs (mean 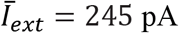). The model average threshold growth lies within the range observed experimentally. However, in the model the threshold is more flexible: it grows very little for small inputs (down-pointing triangles), but very much for large inputs (up-pointing triangles).

Our final model effectively reproduced the decay time constant of the hMCs (Fig. 6E, 539 ±19 ms for the model, and hMCs 570 ±69ms). The initial value for the decay (at the first 50 ms delay between ramps) is highly sensitive to the first (stimulus) ramp intensity, whereas the probe intensity influences the overall trend of the decay (higher intensities shift the curve upwards). This is because the model’s threshold is directly influenced by the injected current, and it seems to grow more rapidly than the experimentally observed threshold, especially for high input currents (as shown in Fig. 6D). The sigmoidal dependence on the injected current in Eq. (1) plays an important role in shaping the threshold decay curve as well. This happens because the probe ramp used to measure the threshold values at subsequent delays, feeds back into the threshold itself, increasing it until a spike is emitted by the membrane voltage running off to infinity. We did not investigate the details of this feedback mechanism because we were interested in a model that captured the average behavior of a hMC rather than the details of the underlying currents.

We plotted the minimum of the AHP versus the threshold of the spike that generated it (*i.e.*, the spike preceding that minimum – see Fig. 6F). These two quantities have a clear correlation in hMCs, motivating the linear reset rule we used in the model. In fact, the parameter *α* = 0.8 (the slope of the reset potential) was directly extracted by fitting the experimental data with a straight line (pooling all spikes from all injected currents together). Notwithstanding, the slope of the model simulation is almost constant at *α*, whereas the observed hMCs slope remains close to 0.8 over a large range of inputs, decreasing drastically for high inputs due to saturation effects in the membrane dynamics that were not included in the model.

We left detailed biophysical data out of the model because we intended to build a model for large-scale simulations of the circuitry involved in spatial learning and navigation. We tested the robustness of the model by varying the parameters that control the threshold dynamics (Fig. 7). All the four threshold-related features that we considered (threshold increase versus spike number and input, latency, and threshold decay) showed robust behavior over variations in parameters. The AHP-threshold correlation was not significantly affected by the parameters in the *θ* and *θ*_*s*_ equations. The parameters that were the most robust were *Δθ*, *τ*_*θ*_ and *V*_*m*_. On the other hand, the model was more sensitive to alterations in the parameters of the *dθ*/*dt* equation, especially *I*_0_. The measured quantities that showed more marked variations with respect to parameter changes are the first spike threshold (Fig. 7A) and the threshold decay (Fig. 7D).

**Figure 7.**
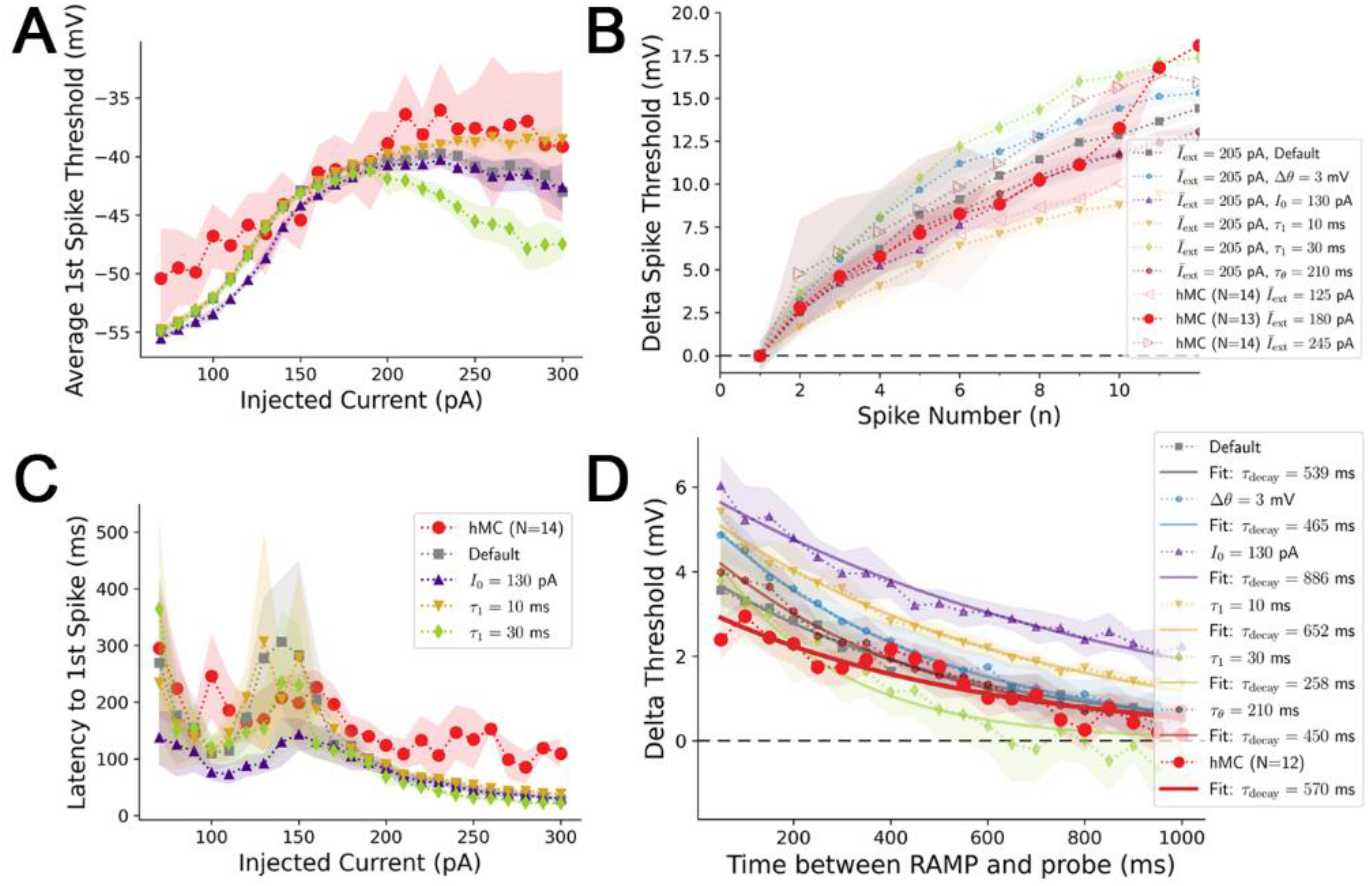
Robustness of the model with respect to threshold parameters. **A.** Threshold as a function of the injected current (same legend as in panel **C**). **B.** Threshold increase as a function of the spike number. **C.** Latency as a function of the injected current. **D.** Threshold decay as a function of the delay after the initial ramp stimulus. The default configuration of parameters is given in Table 1 and shown in Fig. **6**. The hMC data are the same as in Fig. **6E**. The other configurations have the altered parameters given in the legends. The same symbol and color are used for each unique configuration across all panels. Shaded area is the standard error. The model is robust to variations in certain parameters, although choosing the best fit for one of the features may interfere with the fit performance of other features, suggesting a covariance between hMCs membrane parameters. The threshold-AHP minimum covariance plot is not affected by any of these parameters.

The default parameter set for our EIF model that we obtained with our simulations is given in Table 1. It represents an average hMC. Even though our model is robust to variations in the parameters, some changes tend to enhance the fitting to the threshold increase, but worsen the fit to threshold decay, and *vice-versa*. This suggests that the hMCs have correlations between some of their membrane parameters due to correlations of the underlying channel kinetics. Since we were not interested in fitting a complete conductance-based model to the dynamics, we did not look into these correlations in further details. Instead, we simply added a noise term to the membrane equation, generating some variability in the potential simulations and feature statistics of the model behavior, and thus capturing a component of the observed variation in hMCs.

## Discussion

We report herein a biophysical examination of hippocampal hilar neurons that identified at least two salient properties of hMCs that distinguishes them from hilar interneurons. Thus, hMCs displayed: 1) a robust increase in spike threshold with increasing input current that was accompanied by notably delayed spiking, and; 2) strong adaptation mechanisms consisting of a rapidly acting mAHP and a novel spike triggered-increase in threshold acting on a protracted time scale of hundreds of milliseconds. These specific spiking dynamics of hMCs and hilar interneurons organized in a feedforward inhibitory motif converge in constraining the excitability of hMCs in response to synaptic inputs, consistent with their characteristic low firing rates *in vivo* (Goodsmith et al., 2017; Senzai and Buzsaki, 2017). We further developed a cellular model that captures these excitability features and that is thus conducive to examine how the spiking behaviors of hMCs observed *in vivo*, and their resultant computations, emerge from the dynamics of their hilar synaptic inputs.

The spike threshold of hMCs retains a memory of prior spikes (Fig. 4 and 5). Indeed, spike threshold increased steeply over successive spikes making it increasingly difficult to drive spiking (Fig. 4Bi, 4Bii, 4C). A ramping current injection protocol (Trinh et al., 2019) revealed that the spike threshold of hMCs and CA3 pyramidal cells exhibited the greatest threshold increase that recovered the slowest: this emphasizes the similarity between these cell types. The hMC spike-dependent increase in threshold extended for up to 1 s and therefore limits spiking over a timescale that is substantially longer than that of the mAHP. The slowly adapting spike threshold may be caused by the slow recovery from inactivation of Na^+^ channels (Henze and Buzsaki, 2001; Platkiewicz and Brette, 2011; Trinh et al., 2019).

We developed a model that captures salient experimentally-derived excitability properties of hMCs. One important feature of this model is the implementation of a realistic threshold that mimics both the positive feedback characteristics of sodium channel activation and the complex kinetics of its recovery from inactivation. We lack however biophysical insights into the input-dependent threshold of the first spike and its relation to SK channel gating that generate the AHP. The latency to first spike is a consequence of these dynamical effects and more detailed biophysical analyses will be required to better model threshold dependence on natural synaptic input patterns.

The dynamics of our model readily predicts that the substantial and protracted threshold increase following spiking in hMCs impedes high frequency firing. The AHP has the same effect, and augments the increased threshold by its positive covariation with the most recent threshold value (Fig. 6F), *i.e.,* increasing threshold and the concomitant increase in the AHP will act synergistically to limit hMC discharge. These two adaptation mechanisms operate over distinct timescales and are therefore modulating excitability under distinct running average estimates of spike timing. More importantly perhaps, our data could be well fitted without the need for dynamical ingredients that can cause bursting i.e., recurring bouts of high frequency firing. Higher frequencies are seen here at the onset of step currents, as is expected for adaptation processes that are known to preferentially encode changes in input strengths (i.e., their temporal derivative). Our cellular model is therefore now ready for inclusion in a broader network model that will investigate the effects of feedback to GCs, connections between hMC (Ma et al., 2021) and the reciprocal connectivity with hilar interneurons (Larimer and Strowbridge, 2008), and how the emerging network computations are modulated by plasticity (Hashimotodani et al., 2017). This modeling framework will allow to tease out the role of the computations supported by hMCs in the context of pattern separation, completion and comparison (see below).

### Implications for *in vivo* studies of hMCs

While there are far fewer hMCs than GCs (Amaral et al., 1990). they nonetheless occupy a privileged position in the hippocampal network. Mossy fiber collaterals from GC provide a major input to the hilus where they are believed to terminate onto proximal dendritic excrescences of a small number hMCs as well as onto local inhibitory interneurons (Acsady et al., 1998). CA3 pyramidal cells project back to hMC distal dendrites and local interneurons (Scharfman, 1994a; Kneisler and Dingledine, 1995) giving rise to the idea that hMCs acts as comparator of its GC input and its CA3 feedback. The hMCs then feedback to a widespread population of GCs directly (Scharfman et al., 1990; Scharfman, 1995) and via inhibitory interneurons (Scharfman, 1995); i.e., the GC/CA3 pyramidal cell “comparator” now presumably modulates its own input and therefore GC pattern separation and place field responses. From these well described connectivity features, several questions naturally arise, notably on how the putative comparator role of hMC emerges from their excitability dynamics. First, how will a hMC respond to the discharge of GC mossy fiber (Scharfman et al., 1990; Scharfman, 1993) and CA3 pyramidal cells (Scharfman, 1994a, b), separately and in conjunction? Second, what is the effect of hMC spiking on their target GC discharge? The combination of a strong mAHP, dynamic threshold and rapid inhibition suggest that hMC spiking behavior is, as a general feature, powerfully constrained, making unclear the magnitude of their influence on GCs excitability.

Granule cells often fire high frequency bursts in their place field (GoodSmith et al., 2017). Since mossy fibers input to hMCs are strongly facilitating (Scharfman et al., 1990; Lysetskiy et al., 2005; Hedrick et al., 2017; Lituma et al., 2021) and can drive spiking when presented as bursts (Scharfman et al., 1990), hMCs are thus expected to track the bursting of incoming GCs, for instance during spatial navigation. Direct evidence for this supposition comes from paired *in vivo* recordings of GCs and hMCs that shows that the probability of spike transmission rises strongly by the third spike of a burst (Senzai and Buzsaki, 2017). Likewise, the CA3 to hMC synaptic inputs are also facilitating (Scharfman, 1994a; Hedrick et al., 2017). Thus, given the strong facilitatory short-term dynamics of both major excitatory inputs to hMCs, one would expect that a hMC would discharge at high rates as a rodent traversed its place field. This however does not appear to be the case: *in vivo* recordings show hMC firing rates ranging between 8 to 12 Hz in their place field (GoodSmith et al., 2017). The translation of strongly facilitating GC and CA3 pyramidal cell synaptic input to only low frequency hMC discharge may thus be the direct result of the hMC dynamics described herein. The combined actions of a strong mAHP acting over a short time scale and a dynamic threshold acting over a longer time scale are likely to dynamically modulate the encoding features of hMCs in response to synaptic inputs. Strong synaptic inputs are expected to lead to a higher spiking threshold, effectively encoding the onset of GC spike bursts, while filtering out the effects sustained GC spiking. Finally, the lower spike threshold for hilar interneurons compared to hMCs suggests that the prominent inhibitory input to hMCs (Acsady et al., 1998; Larimer and Strowbridge, 2008) will precede or overlap their excitatory input and further suppress their discharge. We hypothesize that these collective dynamics not only contribute to the low firing frequency of hMC discharge observed *in vivo*, but more specifically confers on hMCs the ability to preferentially encode the onset of GC burst spiking during navigation.

### Implications for the induction of plasticity

The hMC to GC synapses are weak and mostly masked by strong feed-forward inhibition (Scharfman, 1995; Hashimotodani et al., 2017). A recent study (Hashimotodani et al., 2017) has shown that hMC projections to GCs, but not to interneurons, undergo strong presynaptic LTP as long as hMC fire multiple bursts (>30) at frequencies >30 Hz). Following potentiation, burst stimulation of hMC axons strongly drive GCs. These results raise a critical question: under what conditions can hMC firing rates increase to the point at which LTP is induced, and they can effectively drive GC discharge? A potential answer comes from experiments suggesting that the septal cholinergic input to hMCs greatly increases their excitability via induction of a spike triggered afterdepolarization (Anderson and Strowbridge, 2014); notably, hilar interneurons are not so affected (Hofmann and Frazier, 2010). The balance of hMC excitation to inhibition would therefore shift from favoring low excitability and inhibition to favoring greater excitability and minimizing inhibition. We suggest that there are two requirements for hMCs to strongly activate their GC targets: firstly, their excitability must be raised by a modulatory input that allows them to discharge at rates >30 Hz and, secondly, their input to GCs must be potentiated. Combined experimental and modeling will be required to investigate this possibility.

Altogether, our results demonstrate that salient features of the *in vivo* behavior of mossy cells can be traced back to the combined dynamics of identified intrinsic properties and GC synaptic inputs. We anticipate that iterative implementations of our experimentally-driven model will allow to delve deeper into how the dentate-hilar network operates during tasks involving spatial learning and navigation that utilizes both path integration and recognition of landmarks (pattern separation).

## Acknowledgements

This work was supported by the Canadian Institutes for Health Research Grant # 153143 to J-C B, AL and LM and by the Brockhouse award funds (493076-2017) to AL and LM and an NSERC award (RGPIN/06204-2014) to AL and a grant from the Krembil Foundation to J-C B, AL and LM. We would like to thank Dr. Kirk Mulatz for his help with technical support and Dr. Érik Harvey-Girard with his help in developing a cutting procedure which would has allowed us to record from the hilar mossy cells *in vitro*.

## Notes

Conflict of interest: The authors declare no competing financial interest.

https://github.com/mgirardis/mossy-cell-dg

### Competing Interest Statement

The authors have declared no competing interest.

